# An efficient hairy root system for genome editing of a β-ODAP pathway gene in *Lathyrus sativus*

**DOI:** 10.1101/2023.04.03.535460

**Authors:** Anjali Verma, Lovenpreet Kaur, Navpreet Kaur, Akanksha Bhardwaj, Ajay K Pandey, Pramod Kaitheri Kandoth

## Abstract

Grass pea (*Lathyrus sativus*) is an ideal legume crop for resource-poor farmers, having resistance to various biotic and abiotic stresses. The seeds of this plant are rich in protein and are the only known dietary source of L-homoarginine. Moreover, it thrives with minimal inputs making it a promising crop in grain legume breeding programs with immense potential for food security. Despite these advantages, the global area under its cultivation has decreased because of the presence of an antinutrient compound, β-N-oxalyl-L-α,β-diamino propionic acid (β-ODAP), which results in neurolathyrism both in humans and animals. Multiple efforts in the past have resulted in the development of improved varieties with low ODAP. Still, due to variations in response to the environment, stable low-ODAP lines have not been developed for large-scale cultivation. In this paper, we report in planta characterization of Oxalyl-CoA Synthetase (OCS) involved in the oxalylating step leading to β-ODAP production. We established a hairy root transformation system for *Lathyrus* and demonstrated the genome editing of *LsOCS*. Further, we show that oxalate accumulates in these hairy roots due to loss-of-function of the *OCS* gene. This is the first report of functional analysis of a *Lathyrus* gene in *Lathyrus*. The hairy root genome editing system we developed can be used as a quick system for functional studies of *Lathyrus* genes.

## Introduction

Grass pea (*Lathyrus sativus*) is a nutrient-rich hardy legume well adapted for growth in arid and semi-arid regions of the world. The legume crop is cultivated on a small scale for food in parts of Middle East, Africa and Asia and for feed in Europe and Australia (Dixit et al., 2016; Vaz Patto et al., 2006). Grass pea has a number of agronomic advantages including tolerance to biotic and abiotic stresses (Campbell, 1997; Gurung et al., 2002; Patto & Rubiales, 2014). It has an intrinsic capacity to fix soil nitrogen and has low input farming requirement than most other crops (Peoples et al., 1995; Kumar et al., 2011). In addition, the seeds are a rich source of protein after soybean; rich in essential amino acids, lysine and threonine (Rotter et al., 1991). It also contains a non-protein amino acid L-homoarginine which is beneficial for cardiovascular health (Rao, 2011). Despite these benefits, the cultivation of grass pea has extensively reduced due to presence of a neurotoxin, β-N-oxalyl-L-α,β-diaminopropionic acid (β-ODAP) in seeds. The compound is believed to cause neurolathyrism, a neurodegenerative disease in humans and animals (Rao et al., 1964).

Multiple research efforts have been undertaken for development of varieties with low or no β-ODAP content. These include germplasm evaluation for identification of low ODAP lines followed by conventional breeding and selection, generation of mutant lines, agronomic management, and development of somaclonal variants (Kumar et al., 2011). Though, these techniques have generated low-toxin varieties, due to variations in response to the environment and genetic factors, stable lines have not been developed for large-scale cultivation (Fikre et al., 2011). Alternatively, tissue culture approach has also been utilized but due to lack of stable transformation protocols (Barik et al., 2005) and recalcitrant nature of *Lathyrus* under in vitro conditions (Barpete et al., 2016), not much progress has been made till date.

The biosynthetic pathway and accumulation of neurotoxin in grass pea is not well understood. The origin of neurotoxin was first hypothesized by oxalylation of α,β-diamino propionic acid (DAPA), which was suggested to arise from asparagine and serine (Murti et al., 1964). Later, this oxalylation was confirmed by partial characterization of first enzyme of this pathway, oxalyl-CoA synthetase (OCS) in crude extracts of *Lathyrus sativus* (Malathi et al., 1970). In a recent advancement, a crystal structure of OCS has been proposed after its identification in *L sativus* (Goldsmith et al., 2022). Further, a heterocyclic amino acid, β-(isoxazolin-5-on-2-yl)-L-alanine (BIA) was identified in *Lathyrus sativus*, *Pisum sativum* and *Lens culinaris* but only in *Lathyrus* it was shown to metabolize into ODAP (Kuo et al., 1998). However, the steps and enzymes involved in conversion of BIA to ODAP remains unclear. Several evidence suggest a role for sulfur metabolism in ODAP synthesis where cysteine synthase (CS) and β-cyanoalanine synthase (CAS) were identified as key regulatory enzymes (Ikegami et al., 1988, 1996; Song et al., 2021). This was also supported by *in silico* characterization of *Ls*CAS where expression analysis under stress conditions have suggested its involvement in regulating ODAP levels (Chakraborty et al., 2018). In addition, nitrogen metabolism also impacts the ODAP levels in plant (Jiao et al., 2006, 2011; Liu et al., 2017) and is regulated by the biochemical activities of CAS (Xu et al., 2017).

The genome of *Lathyrus sativus* has largely been unexplored as the genome sequences were not available until recently (Edwards et al., 2023; Rajarammohan et al., 2023). Only few -omics studies have been carried out to understand the underlying mechanisms of ODAP synthesis (Almeida et al., 2014, 2015; Hao et al., 2017; Verma et al., 2022; Xu et al., 2018; Yang et al., 2014). Hence, identification of key pathway genes with novel functions and engineering them to produce desirable changes using CRISPR/Cas approach was not carried out till date. Enormous progress has been made in sequencing legume genomes which has facilitated development of improved legume varieties for use in pulse breeding programs. In model legume species, *Medicago truncatula* and *Lotus japonicus*, CRISPR/Cas genome editing has been successfully used to understand the function of the key nodulation-associated candidate genes (Curtin et al., 2017) and the symbiotic nitrogen fixation related genes respectively (Wang et al., 2016, 2019). Also, in commercially important legume, *Glycine max* (soybean), hairy root transformation system has been used to carry out targeted mutagenesis by CRISPR/Cas (Jacobs et al., 2015; Sun et al., 2015). Agronomic traits such as seed oil (Do et al., 2019), storage proteins (Li et al., 2019), isoflavone content and resistance to mosaic virus (Zhang et al., 2020) have been improved and mutant soybean lines have also been generated by CRISPR (Bai et al., 2020). Among others, effective transformation protocols and genome editing are being established in *Vigna unguiculata* (cowpea) (Che et al., 2021), *Cicer arietinum* (chickpea) (Badhan et al., 2021) and *Pisum sativum* (pea) (Li et al., 2023). Though, genome editing has been effectively used in legumes to produce desirable agronomic traits, grass pea has most often been neglected due to lack of genomic resources and stable transformation protocols.

*Agrobacterium*-mediated transformation has been the preferred method of choice for genome editing in legumes. In this study, we characterized *LsOCS* functionally by complementing *Arabidopsis AAE3* mutants. Further, we established an efficient hairy root transformation protocol for *Lathyrus* and demonstrated CRISPR/Cas9 gene editing of *LsOCS* in the hairy roots. In addition, we studied the role of *LsOCS* in the biochemical pathway leading to ODAP synthesis in these edited, transgenic, hairy roots. Our study suggests that hairy roots have the potential for use as a quick system for functional analysis of genes in recalcitrant plants, like *Lathyrus*.

## Methods

### 1. Plant materials and growth conditions

*Lathyrus sativus cv*. Pusa-24 seeds were surface sterilized with 70% ethanol for 4 min and washed three times with distilled water. Subsequently, 0.1% HgCl_2_ treatment was given for 10 min and rinsed three times with sterile distilled water. The seeds were germinated on agar plates containing ¼ B5 salts, pH 5.7 (Gamborg, 1970) in growth chamber at 22°C, 60-70% relative humidity, 200 μmol m^-2^ s^-1^ light intensity and 16/8 h photoperiod.

For experiments involving *Arabidopsis,* seeds were sterilized by soaking in 70% ethanol for 3 min and were rinsed with distilled water twice. The seeds were treated with 0.5% sodium hypochlorite for 1 min and were rinsed with sterile distilled water three times. The seeds were plated onto ½ MS medium, pH 5.7 (Murashige and Skoog, 1962) supplemented with 7 g/L agar and 1% sucrose and stratified in dark at 4°C for 2 days. Thereafter, the plates were kept in growth chamber maintained at conditions described above.

### 2. Expression and purification of His-tagged *Ls*OCS recombinant protein

The *LsOCS* expression construct was made by amplifying a full length *LsOCS* ORF by RT-PCR prepared from leaf RNA samples using the primers Ls OCS-F and Ls OCS-R listed in Table S1, and was cloned at the *Nde*І-*Xho*І site of pET28a vector which introduces six histidine residues on the N-terminus of the recombinant protein.

The *Ls*OCS was expressed in *E. coli* BL21(DE3) cells for protein purification. Briefly, a 5 ml culture was grown overnight at 37°C and was used for inoculation of 100 mL Luria Bertani (LB) supplemented with kanamycin (50 mg/L). The protein expression was induced by adding 0.5 mM IPTG to the bacterial culture at OD_600_ value 0.5-0.6. The culture was kept in the shaker at 30°C for 4-5 h and finally, the cells were harvested by centrifugation at 4000 rpm for 10 min at 4°C. The cells were then resuspended in lysis buffer containing 50 mM sodium phosphate, 300 mM NaCl and protein inhibitor cocktail (Promega, Madison, Wisconsin, USA) at pH 7.4. The cells were freeze thawed in liquid nitrogen and ruptured on ice by ultrasonicator (Sonics Vibra Cell, Newton, CT, USA). Cell debris was removed by centrifugation at 13000 rpm for 15 min at 4°C. The supernatant obtained was then loaded onto cobalt resin affinity-chromatography column (Thermo Fisher Scientific, Waltham, MA, USA), equilibrated with pH 7.4 wash buffer containing 50 mM sodium phosphate and 300 mM NaCl. The column was allowed to saturate overnight at 4°C on a rocker shaker. The column was washed with wash buffer containing 10 mM imidazole and finally eluted with elution buffer, pH 7.4 containing 50 mM sodium phosphate, 300 mM NaCl and 150 mM imidazole. The eluate was pooled and concentrated using Vivaspin centrifugal concentrator (Sartorius Stedim Biotech S.A., Aubagne, France) with a molecular weight membrane cut-off of 10kDa. The enzyme preparation was washed thrice with sodium phosphate buffer and finally concentrated in buffer containing 50 mM Tris, 1 mM DTT and 5 mM MgCl_2_, pH 7.5. The molecular weight and purity of the affinity purified *Ls*OCS was ascertained by SDS-PAGE and Coomassie Brilliant Blue R 250 staining.

### 3. Determination of *Ls*OCS activity and kinetics

*Ls*OCS enzyme activity was determined by a coupled enzyme assay (Foster et al., 2012; Ziegler et al., 1987). The assay was initiated with 1 μg recombinant *Ls*OCS in assay buffer containing 0.1M Tris-HCl, pH 8, 2 mM DTT, 5 mM ATP, 10 mM MgCl_2,_ 0.5 mM CoA, 1 mM phosphoenol pyruvate, 0.4 mM NADH, 10 units each of myokinase, pyruvate kinase and lactate dehydrogenase and oxalic acid substrate in a final reaction volume of 100 μl. The reaction mixture was incubated at 37°C for 30 min and rate of NADH oxidation was measured at 340 nm in SpectraMax M5 plate reader (Molecular Devices, San Jose, CA, USA). Oxalate in a range of 5 μM to 400 μM was used to determine K_m_ and V_max._ The experiment was repeated three times independently each having two technical replicates. GraphPad Prism software (GraphPad software, CA, USA) was used to analyze the data.

### 4. Subcellular localization of *Ls*OCS

YFP-*OCS* and *OCS*-GFP translational fusion constructs were prepared by introducing full length *LsOCS-*ORF into pSITE and pMDC83 respectively with primers mentioned in Table S1. Gateway recombination reactions were performed with BP Clonase II and LR clonase II enzyme mixes (Invitrogen, Carlsbad, CA, USA) following manufacturer’s protocol. The *att*B1/*att*B2 sites were added to insert and cloned into entry vector pDONR/Zeo with and without stop codon for N-terminal YFP and C-terminal GFP fusion respectively. The insert was then transferred to destination vector in LR recombination reaction generating YFP-*OCS* and *OCS*-GFP fusion constructs driven by 35S promoter. The plasmids were mobilized into the *A. tumefaciens* strain GV3101. *Agrobacterium* cells carrying the plasmid were inoculated in LB media containing rifampicin (50 mg/L), gentamycin (40 mg/L), and plasmid specific antibiotic for selection, and were incubated overnight at 28°C in incubator shaker. The bacterial culture was centrifuged at 4000 rpm for 10 min and pellet was resuspended in MMA (10 mM MES pH 5.6, 10 mM MgCl_2_, 200 μM acetosyringone) to an O.D._600_ = 0.6. Cultures were incubated for 2-4 h at room temperature in a shaker. The cultures were mixed in 1:1 ratio for co-transformation of GFP/YFP tagged OCS and RFP construct prior to infiltration in *N. benthamiana* leaves (Norkunas et al., 2018) to detect co-localization. After 2-3 days of infiltration, samples were prepared and GFP, RFP and YFP signals were observed under confocal laser microscope (Carl Zeiss, Jena, Germany) with excitation wavelength at 488 nm, 561 nm, 514 nm respectively.

### 5. Functional Complementation in the *Arabidopsis thaliana OCS* (*AAE3*) mutants

The T-DNA insertion lines of *Arabidopsis thaliana ACYL-ACTIVATING ENZYME3 (AAE3)* (At3g48990) were obtained from ABRC at The Ohio State University. To identify homozygous lines of *aae3-1* (SALK_109915), gene specific primers and left border primer of T-DNA insertion line were designed at http://signal.salk.edu/tdnaprimers.2.html. Genomic DNA from rosette leaves was isolated and amplified by PCR using the primers LP, RP and LBa1 listed in the Table S1. Genomic DNA of *Arabidopsis thaliana* ecotype Columbia was used as control. The homozygous lines were selected for bulking of seeds. *Arabidopsis aae3-1* plants were transformed with the *OCS* constructs using *Arabidopsis* floral dip method (Clough & Bent, 1998). *aae3-1* plants were transformed with pRI101 *OCS* having 35S promoter and pRI101 *OCS* with *AAE3* promoter to complement the mutant plants.

### 6. Designing of *LsOCS* CRISPR constucts

The CRISPR/Cas gRNA for *LsOCS* was designed with the help of a web-based tool, CRISPR-direct (https://crispr.dbcls.jp/). As genome information of *Lathyrus* was not available, the gRNA identified were checked for off-targets against transcriptome sequence data (Verma et al., 2022). The binary vector pKSE401 was used for construction of CRISPR construct. The oligonucleotides specific to the target site were annealed and ligated into *Bsa*I site of pKSE401(Xing et al., 2014). Briefly, for designing single target construct, equal volumes of 100 μM complimentary oligonucleotides were mixed in a 50 μl reaction volume containing 1X NEB 3.1 buffer and Milli Q (MQ) water. The primers were annealed at 95°C for 4 min followed by 70°C for 3 min in a thermocycler (Eppendorf, Hamburg, Germany). The reaction was allowed to cool slowly to room temperature. The gRNA was then ligated into *Bsa*I linearized pKSE401 using DNA ligation Kit v.2.0 (TaKaRa, Kusatsu, Shiga, Japan).

The double gRNA construct was made following Xing et al., 2014. Briefly, the two gRNAs were assembled into an expression cassette by PCR using pCBC-DT1T2 plasmid as a template. Two sets of primers (DT1-BsF/DT2-F0 and DT2-R0/DT2-BsR), listed in Table S1 were used in a 50 μl PCR reaction driven by Phusion™ High-Fidelity DNA Polymerase (Thermo Fisher Scientific, Waltham, MA, USA). The PCR conditions were, one cycle of 94°C for 2 min, 30 cycles of 94°C for 15 sec, 60°C for 30 sec and 68°C for 1 min followed by 68°C for 5 min. The PCR product was inserted into *Bsa*I sites of pKSE401 by following golden gate assembly protocol. The reactions were performed in a thermocycler (Eppendorf, Hamburg, Germany) at 37°C for 5 h, followed by one cycle of 50°C for 5 min and 80°C for 10 min.

To produce *Lathyrus sativus* hairy roots, the pKSE401 binary construct were transformed into *Agrobacterium rhizogenes* ARqua 1 strain. *Lathyrus* cotyledons from 7-10 days old seedlings were used for transformation. Briefly, the seed coat was removed and the cotyledons were split open. Simultaneously, a slanting cut was made to detach the cotyledons from the axis, leaving 70% cotyledons for transformation. About 25-30 cotyledons were submerged in *Agrobacterium* suspension (OD_600_ = 0.18-0.2) harbouring a single/double gRNA construct and were subsequently infiltrated at 20-25 mm of Hg pressure for 20 min in a vacuum chamber connected to a vacuum pump. The cotyledons were blot-dried on sterile filter paper and co-cultivated on sterile filter paper wetted with ¼ B5 media for 3 days at 26°C, and 200 μmol m^-2^ s^-1^ photosynthetic photon flux density in the growth chamber. Thereafter, the explants were washed in ¼ B5 salts solution containing timentin (350 mg/L) to remove excess *Agrobacterium*. The explants were then allowed to grow in hairy root induction media containing MS salts with B5 vitamin and antibiotic timentin (250 mg/L). A control was maintained in similar way by transforming the explants with an empty pKSE401 vector. Hairy roots started emerging in 15-20 days. The 1-1.5 cm root tips were cut and subcultured on MS plates supplemented with B5 vitamin having timentin (250 mg/L) and kanamycin (120 mg/L). These hairy roots were subcultured again on kanamycin containing plates for another 4-5 days, and the transgenic roots selected on kanamycin plates were screened by isolating genomic DNA from the hairy roots by 2% CTAB. The genomic DNA was used for PCR amplification using primers flanking the gRNA target sequence (Table S1). The PCR product was gel-purified by GeneJET™ Gel Extraction Kit (Thermo Fisher Scientific, Waltham, MA, USA), and analysed by sequencing for editing. The sequences were then analysed for indels by Inference of CRISPR Editing (ICE) tool (Synthego, Redwood city, CA, USA)

### 7. RNA isolation and qRT analysis

Total RNA was isolated from plant tissue samples using Spectrum™ Plant Total RNA Kit (Sigma-Aldrich, St Louis, MO, USA) followed by DNase digestion with RNase free DNase Set (Qiagen, Hilden, Germany). RNA samples were quantified on Nanodrop (Thermo Fisher Scientific, Waltham, MA, USA) and cDNA was synthesized with 2 μg RNA using PrimeScript™ 1st strand cDNA synthesis kit (TaKaRa, Kusatsu, Shiga, Japan) in a final reaction volume of 20 μl following the manufacturer’s protocol.

Real-time quantitative PCR was performed using gene-specific primers (Table S1) for selected genes on CFX96 Touch™ Real-Time PCR Detection System (Biorad, California, USA). The amplification conditions were: initial denaturation of 95°C for 30 sec, 45 cycles of 95°C for 30 sec, and 60°C for 60 sec. β-tubulin gene in *Lathyrus* hairy roots and *Arabidopsis*’ *TUB2* gene was used as an internal control for normalization of gene expression. Relative gene expression levels were then analyzed by 2^-ΔΔCT^ method (Livak & Schmittgen, 2001). All reactions were performed in triplicates for three biological replicates and averaged.

### 8. Phenotypic analysis of *aae3* mutant and *OCS* complemented seeds

Seed permeability in *Arabidopsis* was determined by incubating 50-100 seeds in 1% (w/v) 2,3,5-triphenyltetrazolium chloride at 30°C in dark for 24 h (Foster et al., 2012). Seeds were rinsed with water and visualized under stereomicroscope (Leica Microsystems, Wetzlar, Germany). Mucilage was observed by staining with ruthenium red. For this, seeds were first immersed in water for 2 h without any disturbance, followed by staining with 0.01% ruthenium red for 1 h (McFarlane et al., 2014). The seeds were rinsed with water and the mucilage layer was observed under stereomicroscope.

To observe morphological differences, dry *Arabidopsis* seeds were mounted on stubs and coated with gold-palladium (80% gold and 20% palladium) in sputter coater (Quorum Technologies Ltd, Laughton, East Sussex, UK) and observed under a field emission scanning electron microscope (Thermo Fisher Scientific Inc., MA, USA).

### 9. Seed germination and weight analysis

Seed germination rate was quantified by plating 100 seeds each of Col-0, *aae3-1*, *35S:OCS* and *AAE3p:OCS* on ½ MS medium, pH 5.7 supplemented with 1% sucrose and 0.7% agar as described previously. The germination was scored on 5 dpg, 8 dpg and 10 dpg by counting seeds with radicle protrusion through seed coat under stereomicroscope. The average germination percentages of three replicates were calculated.

For seed weight analysis, the seed weight was determined by weighing 100 seeds for each genotype using analytical balance (A&D company Ltd., Tokyo, Japan). The average of three independent sampling was done.

### 10. Estimation of ODAP

ODAP was estimated by OPT assay (Rao, 1978, Addis and Narayan, 1994). Lyophilized 5 mg plant samples were extracted with 60% ethanol overnight. The next day, samples were centrifuged at 4°C for 10 min at 13000 rpm to collect clear supernatant. OPT reagent was freshly prepared by dissolving 50 mg o-Phthalaldehyde (Sigma-Aldrich, St Louis, MO, USA) in 500 μl 95% ethanol followed by the addition of 100 μl β-mercaptoethanol and 49.4 ml 0.5 M potassium borate buffer, pH 9.9. For hydrolysis of ODAP, 200 μl 5 N KOH was added to 100 μl extract followed by 100 μl addition of water. The samples were incubated in a dry bath at 100°C for 30 min. Simultaneously, the unhydrolyzed samples were incubated without KOH. The samples were cooled to room temperature for 15 min and 500 μl water was added. 2 ml OPT reagent was added to each sample and incubated at room temperature for 30 min. Thereafter, absorbance for both hydrolyzed and unhydrolyzed samples were determined at 425 nm in SpectraMax M5 plate reader (Molecular Devices, CA, USA). Unhydrolyzed sample readings were subtracted from the hydrolyzed sample values to estimate the ODAP content of samples. Standard curve was prepared as described in Addis and Narayan,1994. The ODAP concentration in the samples was estimated from the standard curve.

### 11. Oxalate estimation

Total oxalate was extracted from fresh tissue samples by following extraction method described by Liu et al., 2009. Briefly, 200 mg fresh tissue samples were ground in 1.6 ml 0.5N HCl in pestle and mortar and transferred to glass tubes. The homogenate was boiled in a water bath for 20 min with mixing the content of the tubes several times during the incubation. After cooling the tubes at room temperature, MQ was added to a final volume of 10 ml. The tubes were kept at 4°C for overnight precipitation of oxalate. Next day, the homogenate was centrifuged at maximum speed for 15 min at 4°C. The clear supernatant obtained was transferred to fresh tubes and pH was adjusted to 7 with 2M KOH. The extract was used for estimation of oxalate with oxalate assay kit (Sigma-Aldrich, St Louis, MO, USA) following the manufacturer’s protocol.

### 12. Statistical analysis

GraphPad Prism software (GraphPad software, CA, USA) was used for statistical analyses. The sample means were compared for significance of difference using one way analysis of variance (ANOVA) followed by Tukey’s multiple comparison test.

## Results

### *LsOCS* complements *AtAAE3*

The lack of genomic information has hampered the characterization of pathway genes in *L. sativus*. As a step towards understanding the pathway genes, we analyzed the transcriptome sequences of *L. sativus* for *Arabidopsis AAE3* homolog and identified *Ls*OCS. The *LsOCS* sequence encoded a protein of 521 amino acids and showed 80.2% identity in the protein BLAST results to *A. thaliana* AAE3. It was also identical to *Medicago truncatula* OCS (90.2%) and *Glycine max* OCS (86.4%), the two leguminous species closer to *L. sativus*.

The *Ls*OCS was expressed in *E. coli* BL21(DE3) cells and was purified to homogeneity (Fig. 1A). Enzyme kinetic studies using oxalate as substrate shows that *Ls*OCS follows Michaelis Menten kinetics with oxalate upto 300 μM followed by substrate inhibition at 400 μM. Furthermore, a K_m_ of 37.7 μM and V_max_ 4.67 μmoles/min/mg protein was determined for the protein (Fig. 1B). These experiments suggested that *Ls*OCS functions as an oxalyl-CoA synthetase in vitro.

**Fig 1:**
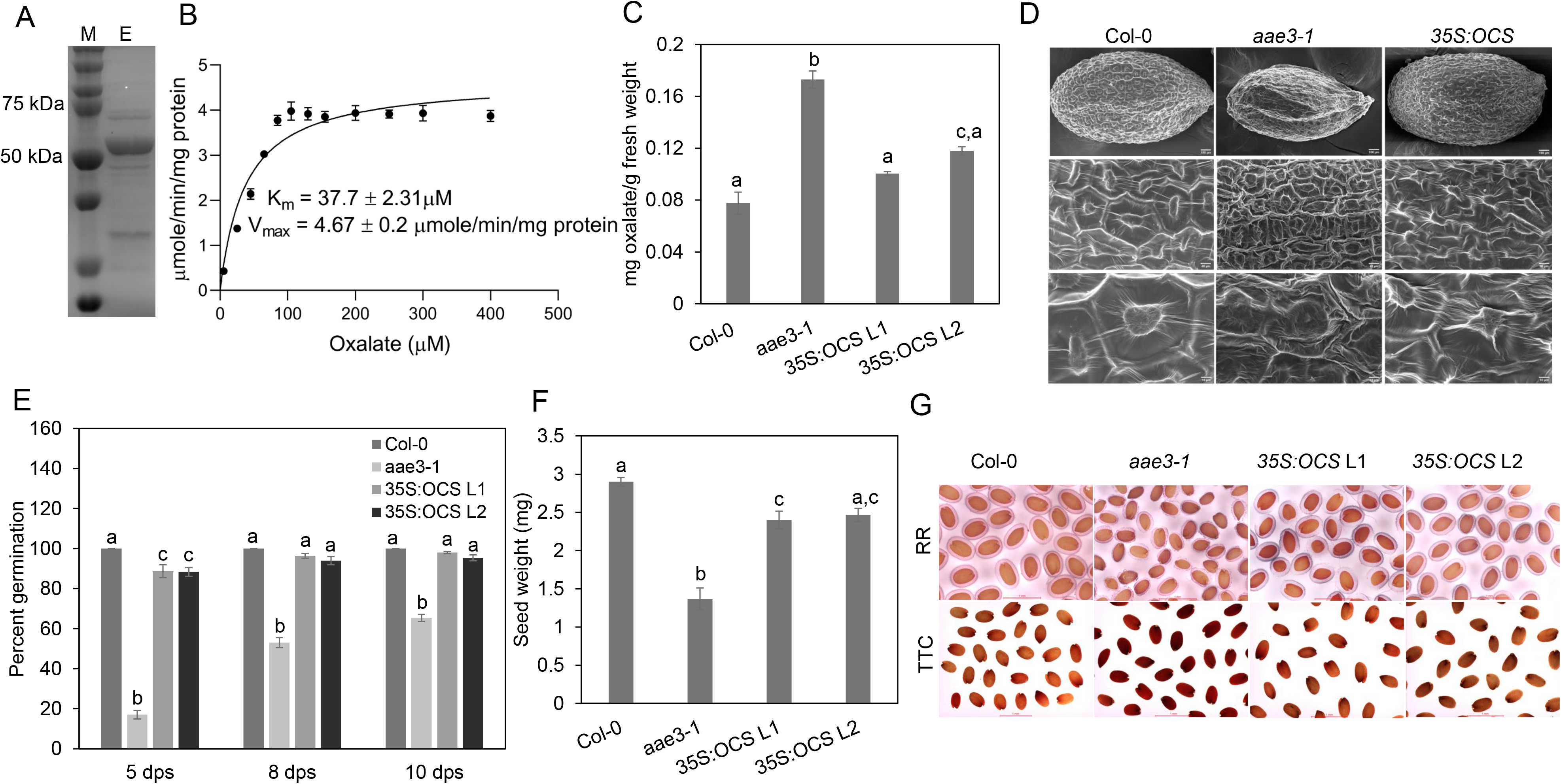
Characterization of *Lathyrus sativus OCS*. A) SDS-PAGE gel image of His-*Ls*OCS recombinant protein purified by cobalt affinity chromatography. Lane M, molecular weight marker, lane E, eluted His tagged *Ls*OCS. B) Enzyme kinetics of *Ls*OCS was performed using oxalate as substrate from 5 μM to 400 μM. K_m_ and V_max_ were calculated using non-linear regression curve fit. Error bars are standard error of means (SEM) of three biological replicates each having two technical replicates. Some data points have too small error bars to be displayed on graph. C) Measurement of seed weight of wild type (Col-0), *aae3-1* and *3SS:OCS* complemented lines, N=100. D) FESEM images of *A. thaliana* seeds. Seed of Col-0 and *35S:OCS* is large in size and is inflated with regular cell wall pattern whereas *aae3-1* seed is small and reduced in size with irregular cell wall pattern in panel A. Magnified images in panels B and C show well-formed columella in seed coat cells of Col-0 and *35S:OCS* seed and collapsed columella in *aae3-1* seed. Bars = 100 μm in panel A, 40 μm in panel B and 10 μm in panel C. E) Seed germination rate of wild type (Col-0), *aae3-1* and *35S:OCS* complemented lines on ½ MS plates. Germinated seeds were counted after 5, 8 and 10 days, N=100. F) Total oxalate content in rosette leaves of *A. thaliana* G) Microscopic images of seeds stained with 0.1% ruthenium red (RR, top panel) and seeds stained with 2,3,5-triphenyltetrazolium chloride (TTC, bottom panel), bars = 1 mm. Error bars in graphs represent standard error of means (SEM) of three biological replicates. Different letters above bars indicate statistically significant differences with Tukey’s multiple comparison test.

Further, in planta functional validation of gene coding for oxalyl-CoA synthetase was achieved by complementing *Arabidopsis aae3-1* plants with *LsOCS* gene. We used the T-DNA insertion lines of *A. thaliana ACYL-ACTIVATING ENZYME3 (AAE3)* (At3g48990), *aae3-1* (Foster et al., 2012) for complementation experiments. The T-DNA insertion was in the first exon of the *AAE3* open reading frame in the mutant (SALK_109915). The plants grown from the mutant line were genotyped by PCR to confirm homozygosity (Fig. S1A). Seeds from the homozygous plants were collected and used in complementation experiments. We visualized the wild and mutant seeds for morphological differences (Fig. S1B and S1C). The mutant seeds appeared small, light in color, and reduced in size compared to wild-type seeds under a stereomicroscope. Also, *aae3-1* showed decreased germination rate, as reported by Foster et al.,2012.

We produced *LsOCS* transgenic lines in the mutant for complementation and verified the *LsOCS* expression (Fig. S1D and S2A). When *aae3-1* plants were complemented with *35S*:*OCS,* the phenotypic differences were restored to wild type, as evident in Fig. 1D, 1E, 1F, and 1G. Loss of *aae3-1* causes oxalate accumulation in the seeds compared to the wild type and leads to the deposition of calcium oxalate crystals in the seed coat (Foster et al., 2012). Decrease in free calcium availability affects cell wall stability, leading to defects in seed coat permeability and mucilage formation (Shi et al., 2017). The seed coat of *Arabidopsis* seeds is characterized by thick-walled hexagonal epidermal cells having volcano-shaped secondary cell walls in the center, columellae (Haughn & Chaudhury, 2005). To observe the rescued phenotype in the complemented seeds, dry seeds were viewed under a field emission scanning electron microscope (FESEM). Mutant *aae3-1* seeds exhibited clear seed coat defects compared to Col-0 seeds. The epidermal cells were irregular and lacked columellae (Fig. 1D). The complemented line seeds had a phenotype similar to wild-type seeds with well-formed seed coat epidermal cells. To visualize the mucilage layer, seeds were stained with ruthenium red and viewed under a stereomicroscope. The mucilage layer of *35S:OCS* complemented *Arabidopsis lines* were comparable to wild-type Col-0 seeds, whereas it was significantly reduced in *aae3-1* seeds (Fig. 1G). In another staining experiment, seeds were incubated in 2,3,5-triphenyltetrazolium chloride to check for the seed coat permeability. The mutant seeds were stained red, indicating defects in the seed coat with increased permeability. However, the wild type and the complemented lines were weakly stained or not stained at all (Fig. 1G). The seed coat defects are attributed to elevated oxalate levels in the plant. Hence, oxalate estimation was done in rosette leaves of 3-week-old *Arabidopsis* plants. The oxalate levels were significantly high in mutant lines. On *LsOCS* complementation, the oxalate levels restored to wild-type levels (Fig. 1C). Further, we measured the germination rate and seed weight of wild-type, mutant, and the *35S:OCS* complemented lines (Fig. 1E and 1F), all comparable to wild-type and drastically different from *aae3-1* seeds. Similarly, with *AAE3p: OCS,* the partial complementation restored some of the phenotypic differences (Fig. S2 B, C, D and E). All these data indicate that *LsOCS* functions as an oxalyl-CoA synthetase in planta.

### Subcellular localization of *Ls*OCS

To predict the subcellular localization of OCS in *L. sativus*, we used WOLFPSORT and TargetP-2.0 tools. Both tools reported conflicting information. WOLFPSORT predicted a predominantly chloroplastic localization, whereas, TargetP predicted a non-organellar location. To confirm whether the protein is localized in cytosol or organelle, we transiently expressed the YFP/GFP fusion constructs of *OCS* in *N. benthamiana* leaves. OCS was tagged with N-terminal YFP, and expression was observed under a confocal microscope. The expression for YFP-OCS was observed in the nucleus as well as in the cytoplasm, which were similar to YFP and RFP controls (Fig. 2A). Further, OCS was also expressed as a C-terminal tagged GFP fused protein to rule out the possibility of N-terminal tag interfering with the plastid localization signals. The expression pattern observed with OCS-GFP was similar to that for YFP-OCS (Fig. 2B). Thus, from the localization pattern, we conclude that the *Ls*OCS is localized in the cytoplasm.

**Fig 2:**
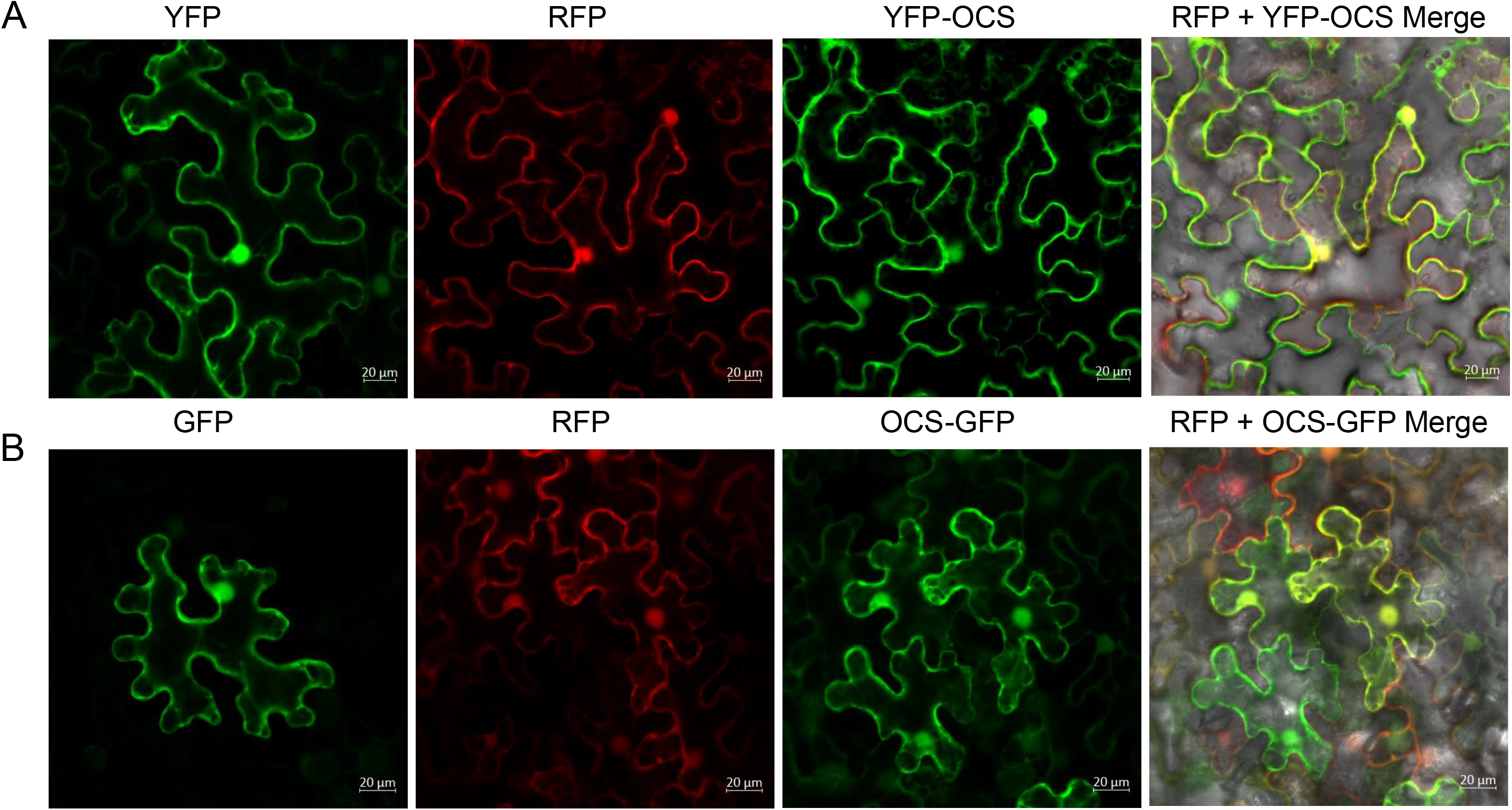
Subcellular Localization of *Ls*OCS protein. A) From left to right, yellow fluorescent protein (YFP) image of epidermal cells of tobacco leaves transiently expressing free YFP, a red fluorescent protein (RFP) image of free RFP, followed by N-terminal tagged YFP-OCS image and a bright field merge of RFP and YFP-OCS. B) From left to right, a green fluorescent protein (GFP) image of free GFP, free RFP image, C terminal tagged OCS-GFP and a bright field merge of RFP and OCS-GFP. Bars = 20 μm.

### CRISPR/Cas9 genome editing in *Lathyrus sativus* hairy roots

For functional analysis of *LsOCS* in *Lathyrus*, we used CRISPR/Cas9 genome editing approach. First, we established hairy root transformation protocols for *Lathyrus*. We used cotyledonary explants of 7-10day old seedlings for *A. rhizogenes* infection. For editing *LsOCS*, we designed a single guide RNA (sgRNA) and cloned it in the vector pKSE401 (Xing et al., 2014) under the control of U6 promoter (Fig. 3A). The *OCS* construct and an empty pKSE401 for use as control, were mobilized into *Agrobacterium rhizogenes* ARqua1 strain to induce hairy roots in cotyledons. Hairy roots emerged after three weeks of infection (Fig S3-A and B). To screen the transformed lines, the roots were cut and subcultured on antibiotic selection plates. Thereafter, PCR was performed on genomic DNA with Cas9-specific primers (Table S1) to identify Cas9 transgenic lines (Fig. 3C). The editing in the transgenic roots were checked by PCR-amplification and sequencing of the target site with primers (Table S1) flanking the target site. As the hairy roots could be chimeric for editing, the sequence of the PCR product was analyzed using Inference of CRISPR Editing (ICE) tool (Synthego, Redwood city, USA) to determine indel percentage. Insertion–deletion or base substitutions around the Cas9 cleavage site, upstream of PAM sequence was identified as editing. For single gRNA, we saw an average editing of 73% in individual roots which suggests that these root cells were chimeric for editing, as confirmed from 15 independent transformed lines. The deletion sizes ranged from 1-11 bases (Fig. 3B).

**Fig 3:**
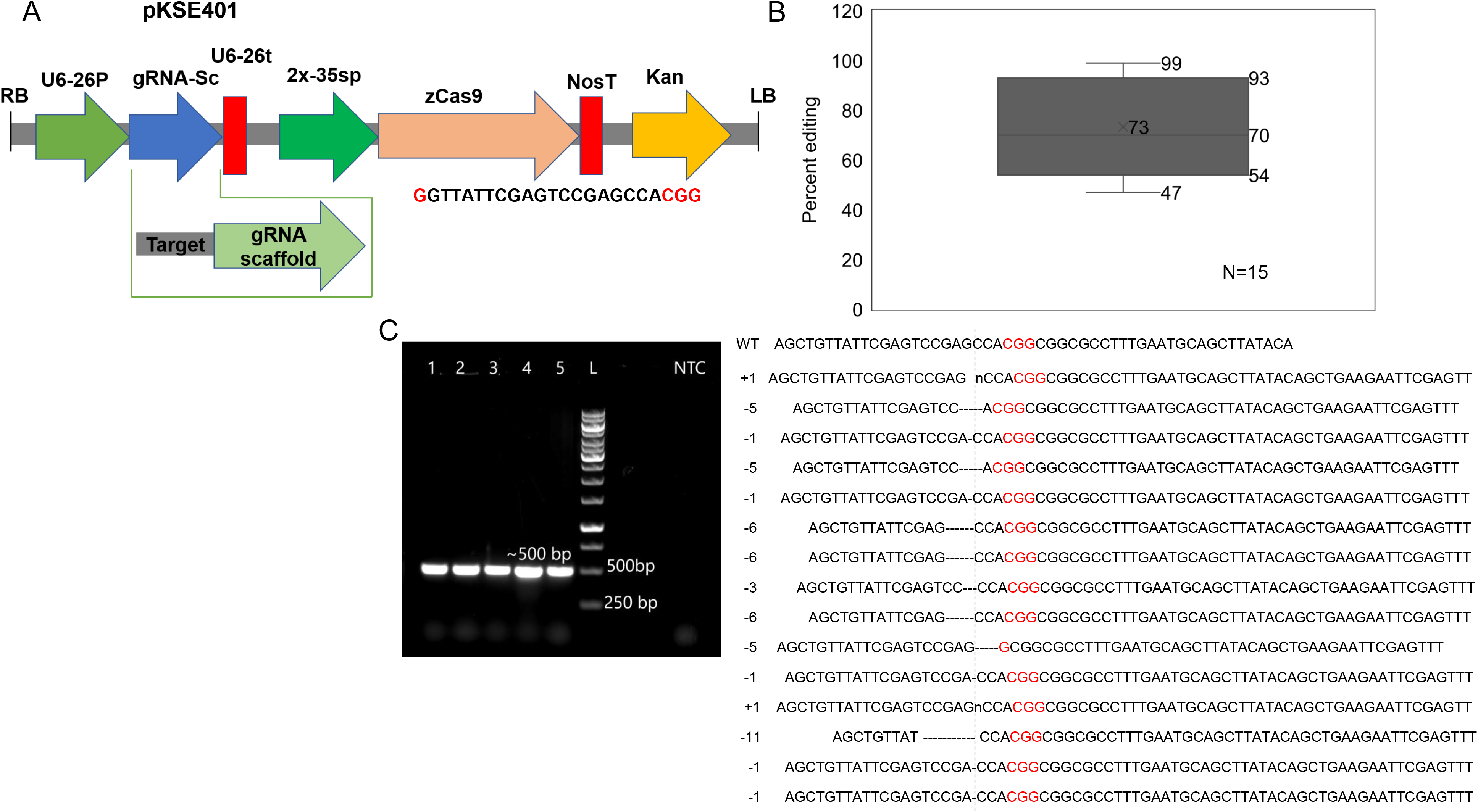
CRISPR/Cas9 genome editing in *Lathyrus sativus* hairy roots using single gRNA. A) Map of pKSE401. B) Box plot of editing percentage in 15 hairy root lines and aligned edited sequences showing PAM sequence (red) next to putative cleavage site (dashed line) C) *Lathyrus* hairy roots after three-week post infection of cotyledons with *Agrobacterium rhizogenes* ARqua1 strain harboring empty vector (control) and D) edited hairy roots (treatment) harboring pKSE with sgRNA for *LsOCS.* E) Screening of transformed hairy root lines on 1% agarose gel after amplification of DNA with Cas9 primers resulting in 500 bp PCR product.

#### Genome Editing with double gRNA construct

In the above method, only sequence analysis of the target site could tell us if the transgenic roots are edited or not. To simplify the procedure, we used a construct with two gRNAs targeting the *LsOCS* gene (double gRNA-dgRNA) (Fig. 4A). The two gRNAs used targeted sites that are 100 bases apart in the gene. In this system, significantly larger deletions can be produced. As expected, small-sized PCR bands compared to the unedited control bands were observed on 1% agarose gels when the genomic DNA was amplified with primers flanking the target sequences in the transgenic hairy root (Fig. 4B). Further, some of these bands were gel-purified and sequenced to examine editing by aligning against the control sequences (Fig 4C). Significant base deletions were observed in most of the roots around the first target site. Thus, the hairy root system provides a rapid, simple, and efficient way for testing the efficiency of CRISPR constructs. The system can be easily employed for understanding the pathway genes and studying the regulatory mechanisms of ODAP production.

**Fig 4:**
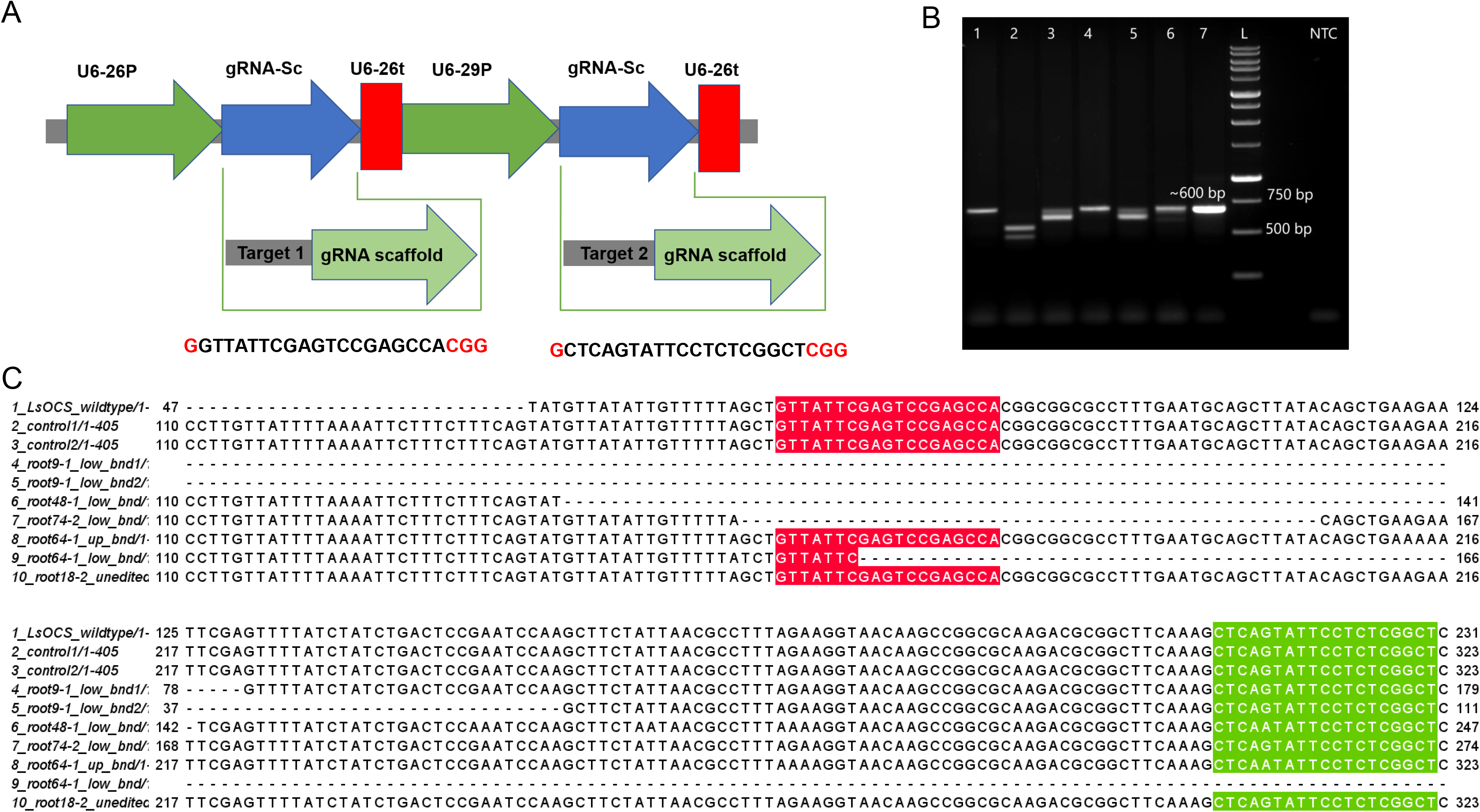
CRISPR/Cas9 genome editing in *Lathyrus sativus* hairy roots using double gRNA. A) dgRNA cassette having two target sites for *LsOCS*. B) Screening of transformed hairy root lines on 1% agarose gel by amplification of genomic DNA with primer set flanking target sites where expected band size of 600 bp confirmed transformed lines. PCR products of small amplicon size with single or double bands (lane 2,3,5,6) confirmed editing in the transformed lines. C) Multiple sequence alignment of nucleotide sequences of *LsOCS* from transformed hairy root lines along with wild type *LsOCS* sequence. Alignment was generated with MUSCLE using Jalview alignment tool. Some sequences were trimmed in 5’ and 3’ region to avoid variable sequencing read length; target site 1 (labelled red) and target site 2 (labelled green).

### Analysis of OCS function in the edited hairy roots

To determine *LsOCS* function in *Lathyrus*, we determined the oxalate content of the *LsOCS* edited hairy roots (Fig. 5). In comparison to the control hairy roots, the edited hairy root lines showed a significant increase in oxalate levels (Fig.5). Further, we observed a good correlation between oxalate levels and the intensity of smaller -sized bands in the PCR products, indicating that roots with higher editing showed more accumulation of oxalate (Fig S3C and S3D). Thus, the data support the role of *Ls*OCS in the degradation of oxalate in the *Lathyrus* plant.

**Fig 5:**
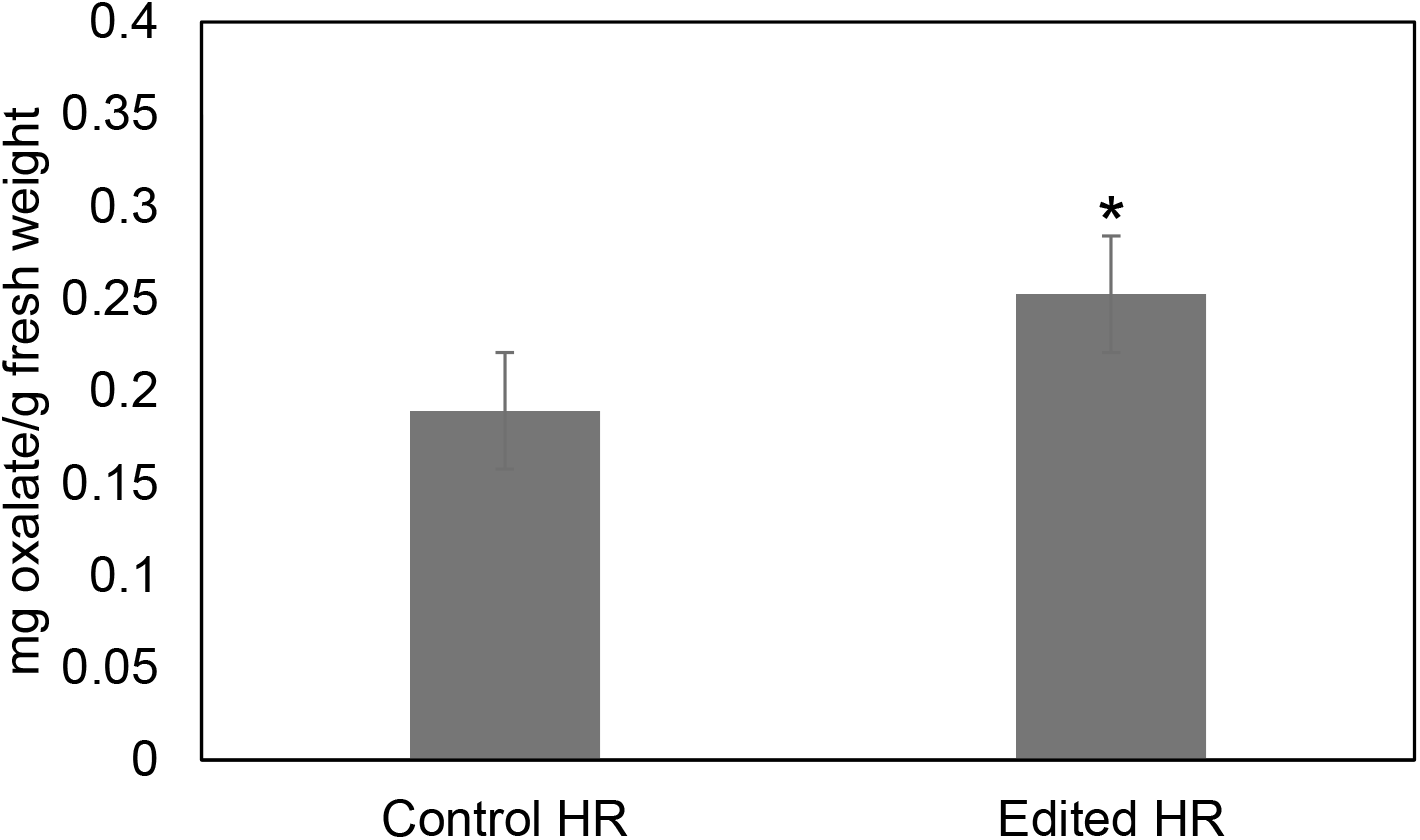
Functional analysis of *Lathyrus sativus* edited hairy roots. Oxalate measurement in three control and five edited hairy root lines, \**p*= 0.0164.

## Discussion

ODAP biosynthetic pathway has been characterized at the biochemical level, however, due to lack of genome information, only recently underlying genes are reported. In this paper, we used *Arabidopsis OCS* mutant *aae3-1* to ascertain the in planta function of *LsOCS,* an enzyme involved in oxalate metabolism and believed to be essential for ODAP biosynthesis. We overexpressed *LsOCS* cDNA to complement the function of the *Arabidopsis aae3* mutant. *Arabidopsis aae3* mutant showed increased oxalate accumulation, resulting in a defect in seed coat formation, reduced seed germination, and reduced seed weight (Foster et al., 2012). The complemented lines showed improved seed germination, seed weight and rectified seed coat defects. They were comparable to wild-type *Arabidopsis* plants (Fig. 1). Moreover, the oxalate accumulated in the mutant plant drastically reduced to levels similar to wild-type *Arabidopsis* upon *LsOCS* complementation. These results show that *LsOCS* functions in oxalate metabolism *in planta*. The in vitro biochemical experiments using a recombinant protein also supported that *Ls*OCS can convert oxalate to oxalyl-CoA (Fig. 1A and 1B).

Further, we did experiments for functional analysis of *LsOCS* in *Lathyrus*. There are no functional analysis tools available in *Lathyrus* to date. Transgenic technology has been tried in *Lathyrus* but with limited success (Barik et al.,2005). The protocols available for genetic transformation in this plant seem genotype-dependent. CRISPR technology for genome editing has been used in multiple crops, including legumes, but its application in *Lathyrus* is yet to be reported. In this work, we established a protocol for hairy root transformation for testing CRISPR technology in *Lathyrus*. The protocols for *Lathyrus* were developed in line with that developed for soybean. We found that within a month we were able to develop transgenic hairy roots in *Lathyrus*. For functional analysis of *LsOCS*, we used two different constructs to edit the *LsOCS* gene: first, we used a single gRNA that targeted the *LsOCS* exon closer to the 5’-end of the gene. PCR amplification of the target site and sequencing of the amplicon confirmed a high editing rate in hairy root cells. Hairy root editing is chimeric. The ICE sequence analysis showed that, on average, 73% of the sequences from individual hairy roots showed deletions (Fig 3B).

To further improve the technology, we used a double gRNA approach, where two gRNAs target the *LsOCS* sequences around 100-150 bp apart. After antibiotic marker-based selection of the transgenic hairy roots, we used a PCR screen to identify edited hairy roots. We were able to detect smaller-sized PCR products in the edited roots (Fig. 4B). Most of the time, the editing was incomplete, indicating the chimeric nature of these roots. However, in some hairy roots, we obtained more than 90% edited sequences. The double gRNA approach is less time-consuming and is helpful for faster detection of edited hairy roots that requires no sequencing confirmation.

As a first step, we measured the oxalate content for functional analysis of *LsOCS* in these edited roots. We found that these roots have increased oxalate levels, as seen in *Arabidopsis* mutants. Furthermore, we found an excellent correlation between the percentage editing in the chimeric roots and the oxalate levels (Fig. S3C and S3D). In highly edited roots, the oxalate accumulation was much higher. These results suggest *LsOCS* functions in the oxalate degradation pathway in *Lathyrus*. The next step will be to test its role in the ODAP pathway, which is more demanding since it requires more tissue owing to lower ODAP levels in the roots.

In conclusion, our experiments demonstrate that *LsOCS* functions as an oxalyl-CoA synthetase *in planta*. We developed a hairy root genome editing protocol and demonstrated the function of *LsOCS* in *Lathyrus*. This is the first report of a gene function study in *Lathyrus* plant and hairy roots could be used as a quick system for gene function studies in this recalcitrant crop.

## Supporting information

Genotyping of mutants and expression analysis

Analysis of pAtAAE3 driven LsOCS plants

Functional analysis of edited hairy roots

## ACKNOWLEDGEMENTS

This work was supported by DST-SERB Core Research Grant No. CRG001736 to PKK. PKK acknowledges Ramalingaswami Fellowship from Department of Biotechnology, Government of India. AV is a recipient of CSIR-UGC fellowship for research. DelCON, Gurgaon, India is acknowledged for access to online journals.

## Figure Legends

**Fig S1:** Microscopic image of wild and mutant seeds showing morphological differences. A) Genotyping of *AAE3* mutants. Identification of homozygous mutants of *aae3-1* by amplification of genomic DNA using gene specific primers (LP and RP, expected band size 1100 bp) and T-DNA left border primer and gene specific right primer (LBa1 with RP, expected band size smaller than WT in lane 2). B) Wild type Col-0 seeds C) *aae3-1* seeds. D) Relative expression of *AAE3* and OCS in rosette leaves of 3 week old seedlings of wild type (Col-0), *AAE3* mutant (*aae3-1*) and *AAE3* mutant complemented with *LsOCS* (*35S:OCS* L1 and *35S:OCS* L2). Error bars in graphs represent standard error of means (SEM) of three biological replicates. Different letters above bars indicate statistically significant differences with Tukey’s multiple comparison test.

**Fig S2:** Functional analysis by complementation using *AtAAE3* promoter driven *LsOCS* plant binary construct. A) Relative expression of *AAE3* and OCS in rosette leaves of 3 week old seedlings of wild type (Col-0), *AAE3* mutant (*aae3-1*) and *AAE3* mutant complemented with *AAE3p:OCS*(*AAE3p:OCS* L1 and *AAE3p:OCS* L2). Error bars in graphs represent standard error of means (SEM) of two biological replicates. Different letters above bars indicate statistically significant differences with Tukey’s multiple comparison test. B) Measurement of seed weight of wild type (Col-0), *aae3-1* and *AAE3p:OCS* complemented lines, N=100. D) FESEM images of *A. thaliana* seeds. Seed of Col-0 and *AAE3p:OCS* is large in size and is inflated with regular cell wall pattern whereas *aae3-1* seed is small and reduced in size with irregular cell wall pattern in panel A. Magnified images in panels B and C show well-formed columella in seed coat cells of Col-0 and in *AAE3p:OCS* seed and collapsed columella in *aae3-1* seed. Bars = 100 μm in panel A, 40 μm in panel B and 10 μm in panel C D) Seed germination rate of wild type (Col-0), *aae3-1* and *AAE3p:OCS* complemented lines on ½ MS plates. Germinated seeds were counted after 5, 8 and 10 days, N=100. E) Microscopic images of seeds stained with 0.1% ruthenium red (RR, top panel) and seeds stained with 2,3,5-triphenyltetrazolium chloride (TTC, bottom panel), bars = 1 mm. Error bars in graphs represent standard error of means (SEM) of three biological replicates. Different letters above bars indicate statistically significant differences with Tukey’s multiple comparison test.

**Fig S3.** *Agrobacterium rhizogenes* mediated hairy root transformation in *Lathyrus sativus*. A) *Lathyrus sativus* seeds. B) Seeds plated for germination on ¼ B5 media plates after sterilization. C) Seedling growth after 7 days post germination. D) Excised cotyledons dipped in *A. rhizogenes* suspension harboring desired CRISPR construct. E) Agroinfiltration of cotyledonary explants under vacuum created in a desiccator. F) Co-cultivation of infiltrated explants on sterile filter paper soaked in ¼ B5 meida. G) Cotyledonary explants kept on hairy root induction media post co-cultivation with cut surfaces. H) Emergence of hairy roots from cut surfaces of cotyledons after 15-20 days of growth in hairy root induction media. I) Excised hairy roots kept on selection media for screening of positive transformants. J) Selected transformed hairy root biomass on growth media plate.

## Notes

### Competing Interest Statement

The authors have declared no competing interest.

