## Supplementary figures and images for "An efficient hairy root system for genome editing of a β-ODAP pathway gene in *Lathyrus sativus*"

### Analysis of pAtAAE3 driven LsOCS plants

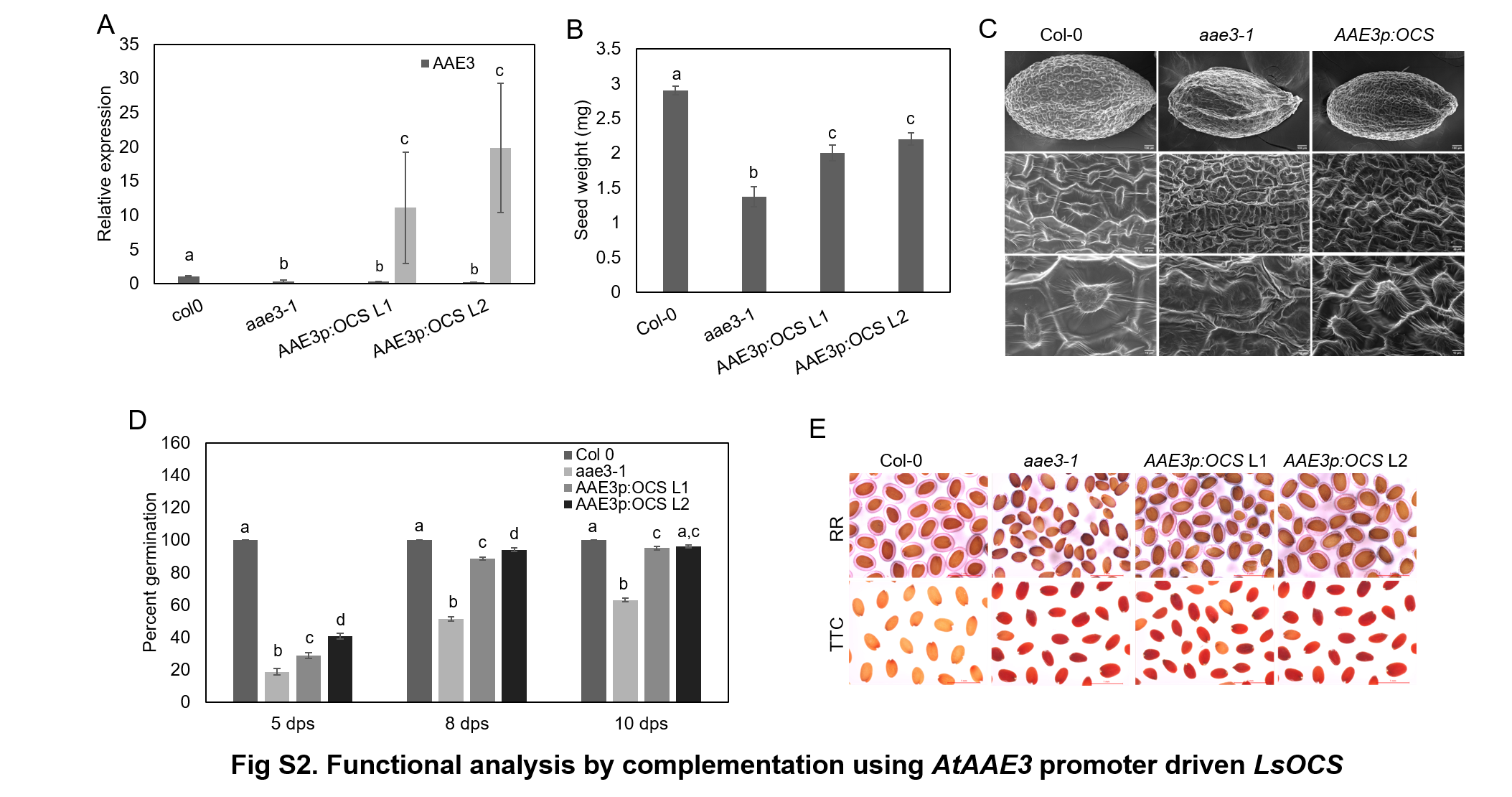

### Functional analysis of edited hairy roots

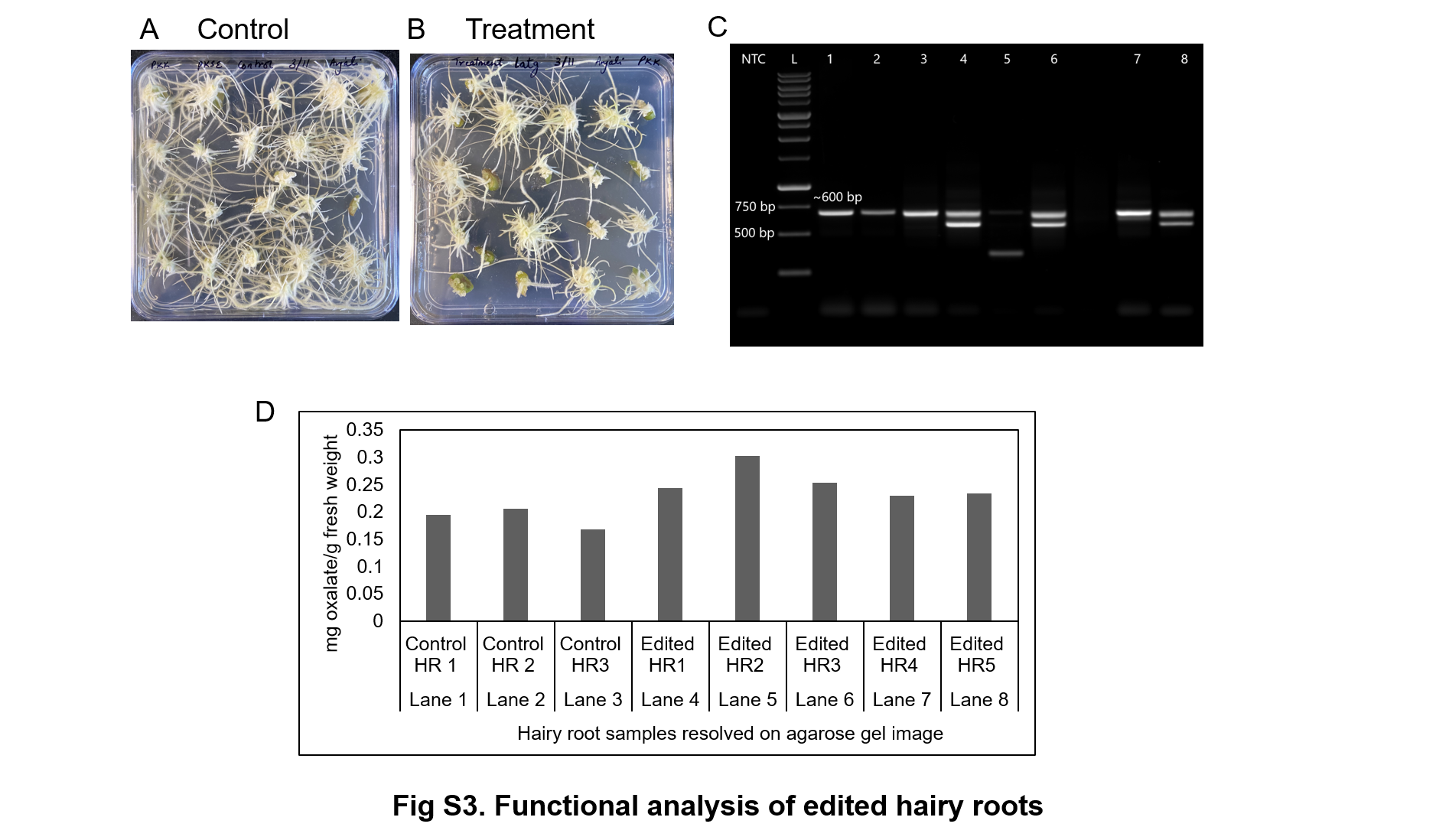

### Genotyping of mutants and expression analysis

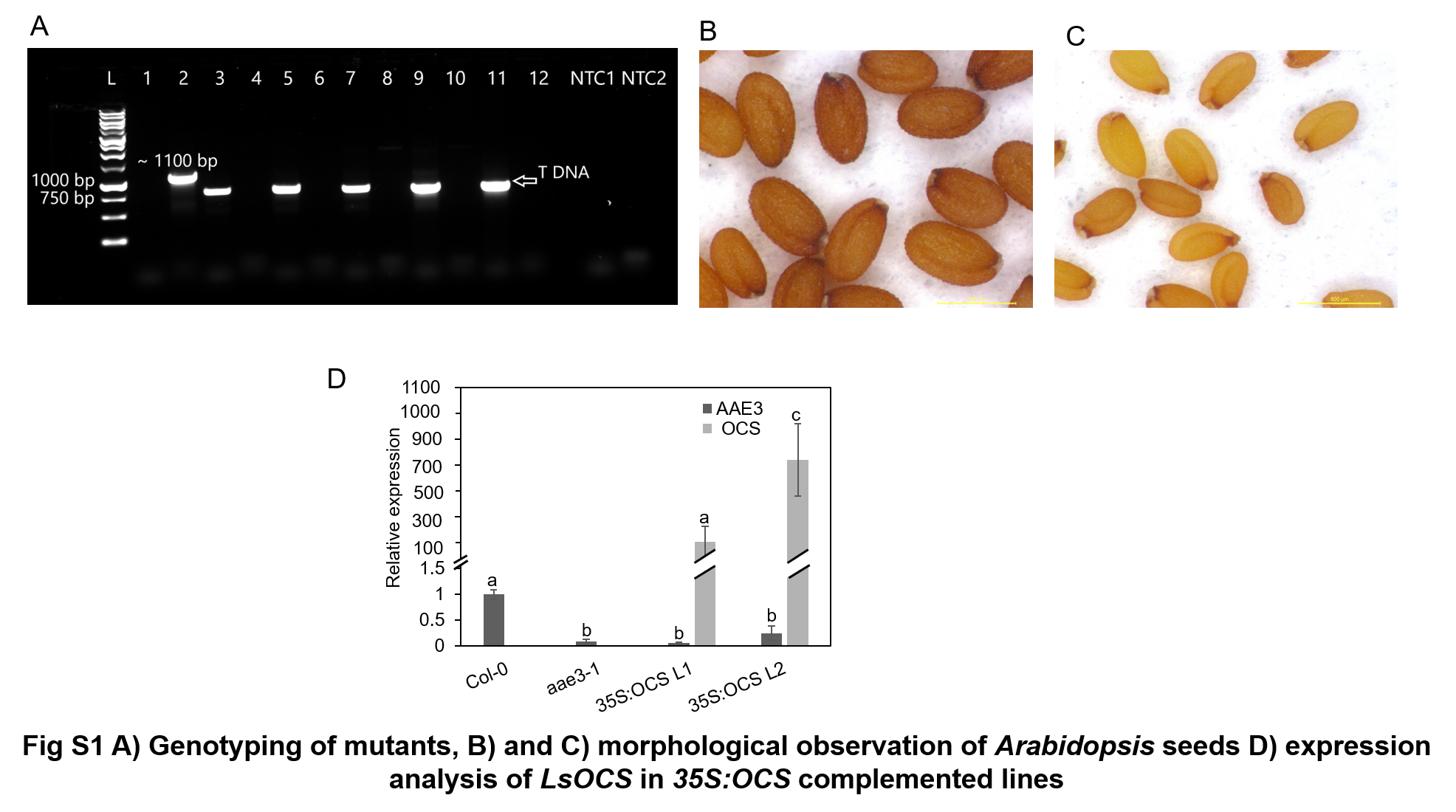
