## Supplementary material for "An efficient hairy root system for genome editing of a β-ODAP pathway gene in *Lathyrus sativus*": Table S1

| **Table S1** | |
|---|---|
| **Primers for funding His tag to LsOCS protein** | |
| Ls OCS-F | GGAATTCCATATGGAAACCGCAACCACCCTCAC |
| LsOCS-R | CCGCTCGAGTCAAACTTTAGAAACAAAGTGTTCTGCTAC |
| **Primers for localization study** | |
| OCS BP Fw | GGGGACAAGTTTGTACAAAAAAGCAGGCTTCATGGAAACCGCAACCACCC |
| OCS BP Rv | GGGGACCACTTTGTACAAGAAAGCTGGGTCTCAAACTTTAGAAACAAAGTGTTCTGC |
| OCS-Ctr BP Rev | GGGGACCACTTTGTACAAGAAAGCTGGGTCAACTTTAGAAACAAAGTGTTCTGC |
| **Primers for genotyping** | |
| SALK 109915-LP | TCAAAGACACATGTGCAAAGG |
| SALK 109915-RP | GGATAGTACAAGGTCCGAGCC |
| LBa1 | TGGTTCACGTAGTGGGCCATCG |
| **Primers for complementation study** | |
| OASComp-F | GGAATTCCATATGGAAACCGCAACCACCCTCAC |
| OASComp-R | CCGGGATCCTCAAACTTTAGAAACAAAGTGTTCTGCTAC |
| AtOAS promoter-F | GCTAACCTGCAGGATGAACCCACCAATATAAGTATAAG |
| AtOAS promoter-R | GGAATTCCATATG GGTTACGTCGGAGAGATAAACACGAGAG |
| **Primers for creating sgRNA construct** | |
| Lathyrus OASgRNA1-F | GGGCGTTATTCGAGTCCGAGCCA |
| Lathyrus OASgRNA1-R | AAACTGGCTCGGACTCGAATAAC |
| **Primers for creating dgRNA construct** | |
| DT1-OCS1 F | ATATATGGTCTCGATTGGTTATTCGAGTCCGAGCCAGTT |
| DT1-OCS1F0 | TGGTTATTCGAGTCCGAGCCAGTTTTAGAGCTAGAAATAGC |

| | |
|---|---|
| DT2-<br>OCS2R0 | AACAGCCGAGAGGAATACTGAGCAATCTCTTAGTCGACTCTAC |
| DT2- OCS2<br>R | ATTATTGGTCTCGAAACAGCCGAGAGGAATACTGAGCAA |
| **Primers for transgene check** | |
| Cas9 F | |
| Cas9 R | |
| **Flanking primers for amplifying gRNA target sites** | |
| fgRNA-<br>OAS-F | CTCACTCTCATCTTAACGAATTAG |
| OASseq-R4 | ACCCAGTGAACTCAGCAATC |
| **Primers used for qPCR** | |
| qOCS F | CGCCTTTGAATGCAGCTTATAC |
| qOCS R | GCCGGCTTGTTACCTTCTAA |
| qLsTub F | TGCCTAGGATCAGCAGCACA |
| qLsTub R | TCAGTGTCCCTGAGCTCACT |
| qAAE3 F | TGGGGAAGAGATTAACTGTGC |
| qAAE3 R | GCGCTGAATCTTACCAGAGG |
| qAtTUB2F | ACTGTCTCCAAGGGTTCCAGGTTT |
| qAtTUB2R | ACCGAGAAGGTAAGCATCATGCGA |
